# Graph theory for the analysis of micro-electrode array recordings of human brain slices – framework and benchmarking

**DOI:** 10.64898/2026.08.10.743867

**Authors:** Jonas Ort, Victoria Sofie Witzig, Aniella Bak, Jan Heckelmann, Ann-Kathrin Roeb, Hussam Hamou, Anke Höllig, Yvonne Weber, Hans Clusmann, Daniel Delev, Henner Koch

## Abstract

Micro-electrode array (MEA) recordings are widely used to characterize functional connectivity in neural cultures and have gained traction for the analysis of human brain slices. However, the impact of graph construction methodology on the resulting network topology has not been systematically quantified. Here, we benchmark three methods - shared spiking activity, Pearson cross-correlation, and the spike time tiling coefficient (STTC) - across 37 recordings from human cortical slice cultures classified into low, moderate, and high activity groups. We show that method choice alone produces large topological differences (Cohen’s d = 0.86–1.14 for clustering coefficient, d > 1.0 for node count), while higher-order features such as modularity remain stable. Each method exhibits a distinct sensitivity profile: shared spiking detects activity-dependent changes primarily through network size, correlation uniquely captures clustering differences, and STTC combines strong biological sensitivity with negligible parameter dependence across lag windows (all d < 0.1). Within shared spiking, z-score normalization dominates all other parameter choices (d > 1.0 versus bin size effects of d < 0.23), functioning as an implicit analytical null model that fundamentally reshapes the edge set rather than merely rescaling weights. Inter-method edge overlap is low (Jaccard index 0.08–0.45) and activity dependent, demonstrating that these methods identify substantially different connections from identical data. Our results reveal that methodological choices including construction method, threshold, and normalization introduce hidden degrees of freedom with effect sizes comparable to the biological signals being measured. We provide practical recommendations for parameter selection, reporting, and cross-method validation in MEA-based network neuroscience.

**Author Summary:** When we record electrical activity from brain tissue using grids of electrodes, we can ask how different sites influence one another and map the tissue as a network of connections. Thanks to novel culturing methods, this approach is increasingly used to study human brain slices. However, deciding what is “connected” is not well defined. Researchers use several different methods, and it has never been clear how much this choice shapes the network they end up describing. Here we compared three widely used methods on 37 recordings from human cortical slices spanning a range of activity levels. We found that the method alone can change the apparent structure of the network as much as real biological differences do. The methods frequently disagreed about which connections exist and some technical choices, including normalization techniques, had surprisingly large effects. Because these hidden choices can rival the biological signal, we provide this benchmarking work with practical recommendations for selecting, reporting, and cross-checking methods, so that network studies of brain tissue become more transparent, comparable, and reproducible.

## Introduction

Micro-electrode array (MEA) recordings offer unique insights into spatial electrophysiology on a meso-scale and have become an essential research method in modern neuroscience. Ranging from stem-cell-culture experiments to cultivated slice culture models, MEAs are used to investigate neuron-population by observing the essential biological readout of the neurons: the action potential (AP). MEA data offers insights into the complex interplay of biological neural networks. However, the data analysis is challenging due to the spatial and temporal components, the data volume, and the complexity of describing and differentiating biologically meaningful activity from random process in the observed networks.

New culturing methods can be used for the investigation of human brain tissue in-vitro and offer a novel model for investigating biological and pathological mechanisms ranging from epilepsy to neuro-oncological research questions [1,2]. Due to the multidimensionality of the data, there are still various open questions about best practices in all subfields of the analysis ranging from spike detection, spike sorting, to network analysis [3–5]. The identification of robust and reoccurring neuronal spike patterns is mathematically and computationally complex [6–8]. These challenges in part explain why a significant amount of publications that use MEA data of human (and rodent) brain slices employ simplistic measurement parameters such as firing rate, number of active channels, or number of bursts. While these parameters are physiologically meaningful, they come short in describing the underlying networks. In other neuroscientific research methods, Graph Theory has been widely applied as a tool for characterizing networks with the use ranging from fMRI studies to EEG or MEG data [9–16]. Graph Theory itself is a form of pure mathematics that originated from Leonhard Eulers solution to a topological puzzle and describes, in its essence, geometrical relations of objects [17,18]. From the 1960s onward, it has found several real-world applications in its data-driven alter ego of network science, which has been applied to a variety of fields including, social science or transportation [19–25].

In short, a graph consists of two building blocks, the nodes (or vertices) and edges (links between nodes). These basic elements can be used to abstractly represent components of any real-world networks, making the method applicable in a wide range of disciplines.

A graph object’s properties are essentially defined by its structure and a multitude of parameters and algorithms have been established that can be used to characterize graphs.

Nodes are connected via edges and the amount of edges a node has is the degree of that node. Further, edges can be described with a weight that can be used to exhibit the strength of the respective connection. Networks can further be analyzed including their degree distributions, connectedness, efficiency, scale-free properties, or small-world properties.

While graph theory’s versatility represents a significant analytical strength, this flexibility simultaneously poses a fundamental challenge: the critical dependence on input parameter selection and graph construction methodology. Both can fundamentally change graph topology and research conclusion. Published MEA analysis that leverage graph theory do suffer from lacking standardization. Further, connections and networks in graphs do not equal meaningful neurobiological processes but are a mere representation of possible underlying structures. This is pivotal also considering the heterogeneity of MEA recording quality and the variability between slices – even more so in human samples. The use of graph theory for MEA-recordings itself is not novel, however, often methodological papers exploring this is based on simulated data.[26]

In this publication we investigate the application of graph theory on human brain slice MEA recordings focusing on the impact of construction methodology and systematic benchmarking of construction parameters. Specifically, we benchmark three graph construction methods against one another on identical recordings: **shared spiking**, which discretizes spike trains into time bins and links electrodes co-active within the same bin; **correlation-based construction**, which computes pairwise Pearson correlations between binned firing rate vectors; and the **spike time tiling coefficient (STTC)**, which operates directly on spike times without binning (Fig 1). Each method was applied to 37 multi-electrode array recordings from human cortical slice cultures spanning a broad range of activity states, and evaluated along three axes: inter-method divergence, sensitivity to biological activity level, and sensitivity to construction parameters.

**Fig 1.**
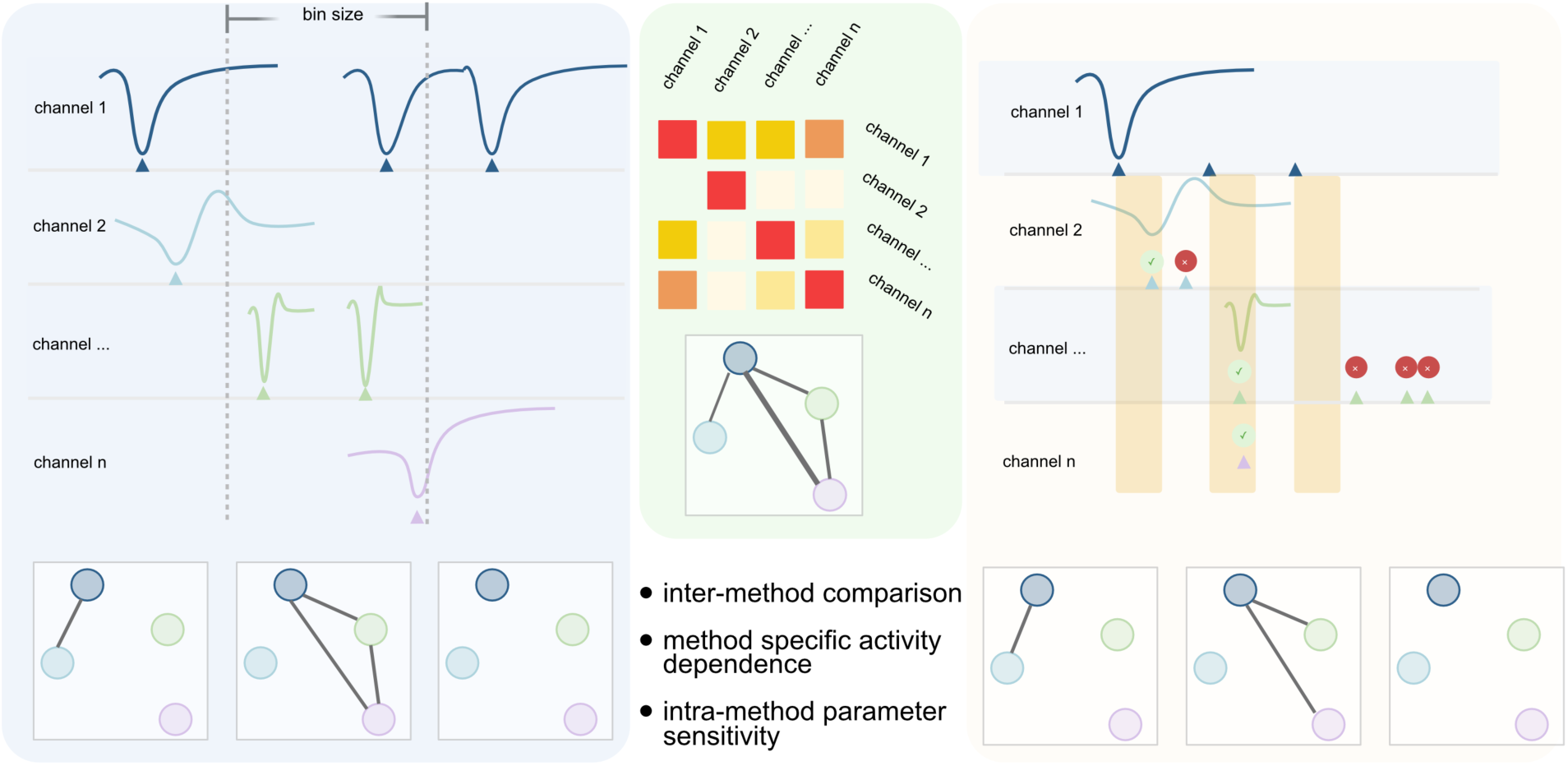
Graph construction methods for functional connectivity analysis. Three approaches were applied to multi-electrode array recordings from human brain slice cultures. (Left) Shared spiking discretizes spike trains into time bins and connects electrodes with simultaneous activity; edge weights reflect cumulative co-active bins across the recording. (Centre) Correlation-based graph construction computes pairwise Pearson correlation between binned firing rate vectors across all electrodes; threshold passing correlations are retained as weighted edges. (Right) The spike time tiling coefficient (STTC) places symmetric time windows (±Δt) around each spike and quantifies the proportion of partner spikes falling within these windows, corrected for total time covered. Unlike the other methods, STTC operates directly on spike times without binning. All three methods produce weighted, undirected graphs compared across inter-method divergence, activity-level sensitivity, and parameter sensitivity.

## Results

MEA recordings from human brain slice cultures varied substantially in both overall activity and activity mode, ranging from sparse, spatially disperse firing to rhythmic, synchronized population events (Fig 2A–C). To ensure that method comparisons were not confounded by this heterogeneity, all 37 recordings were assigned to activity groups. Initial assignment by visual inspection of raster plots was validated by unsupervised clustering across eight quantitative activity features, which identified mean firing rate and network burst frequency as the primary discriminators of activity state (Fig 2D; see Methods). Applying the resulting data-driven cutoffs yielded three groups of comparable size, low (n = 13), moderate (n = 14), and high synchronized activity (n = 10), which form the basis for all subsequent analysis (see Fig 2).

**Figure 2.**
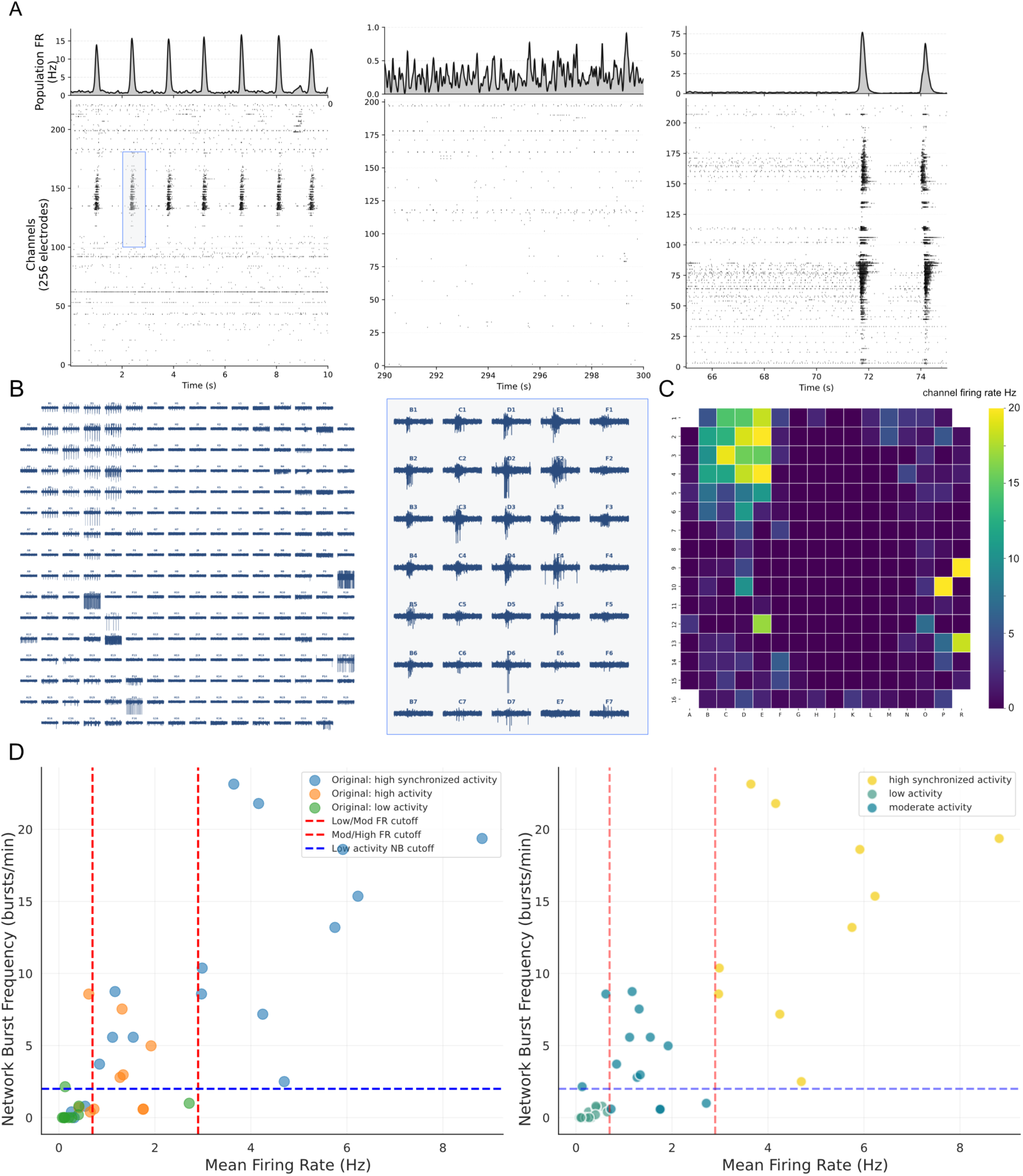
(A) MEA recordings of human brain slices show remarkable variation in general activity and activity modes. The 10-second excerpts of three different human-brain slices with a rhythmic synchronized activity (left), sparse low activity (middle), and a two-mode situation with a relatively low activity and two sudden huge network activations (right). (B) Signal traces of a two second snippet of the recording depicted in (A) with zoomed in activity of the network activation in (A) (small blue rectangle). MEA recordings offer unique spatio-temporal insights into brain slice activity. (C) The mean firing rate for the full recording shows an active cluster in the upper left area of the recordings electrode grid which also corresponds with the short snippet of traces in (B). ***(E)*** Recordings used for this publication where screened from the rasterplots with visual different activity modes between three main broad patterns: high activity with visually synchronized population activity, moderate activity predominantly lacking population synchronization, and low disperse activity. From the data, a kmeans-clustering was performed to ensure the validity of three distinct groups that would serve as data-basis for graph method comparison (see methods for details). The mean firing rate (Hz) and network burst frequency (burst/minute) were identified as the most important features. D (left) shows the original visual assignment and D (right) the reassignment after applying the clustering-derived thresholds. These three activity groups were used for the further benchmarking in this study.

**Fig 3.**
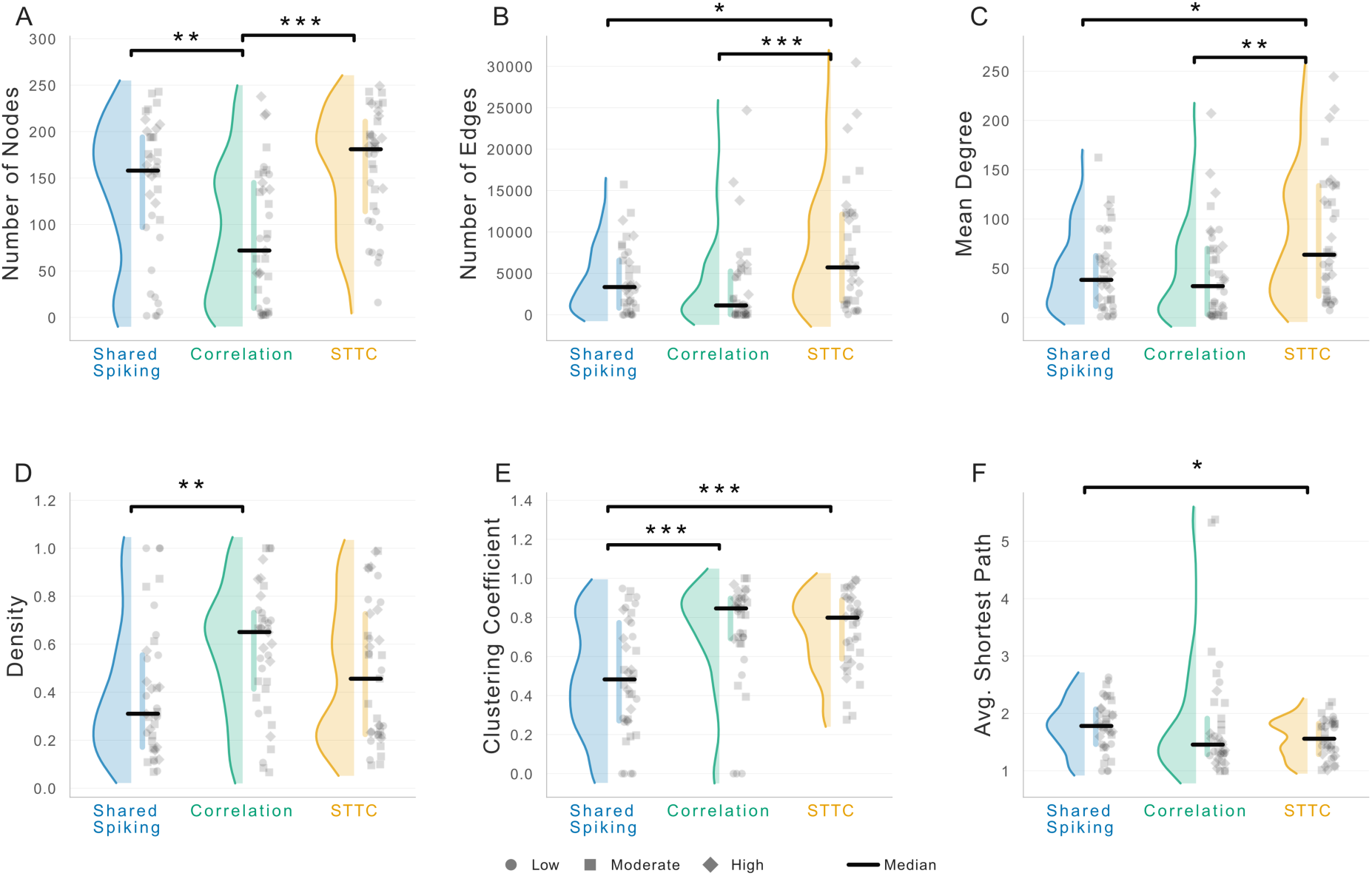
Cross-method comparison of network topology metrics. Half-violin plots show the distribution of six graph-theoretic metrics across all 37 recordings for each graph construction method (shared spiking, blue; correlation, green; STTC, orange). Individual recordings are shown as jittered points with shape encoding activity level (circles = low, squares = moderate, diamonds = high). Black horizontal lines indicate group medians. Significance brackets show pairwise Mann–Whitney U tests (Bonferroni-corrected); only significant comparisons are displayed. (A) Number of nodes retained in the largest connected component. (B) Total number of edges. (C) Mean node degree. (D) Network density. (E) Weighted clustering coefficient. (F) Average shortest path length (inverse-weight).

### 1. Different Graph Constructions Methods yield structurally different networks

To examine the influence of graph construction method on the network topology, we compared the three methods (SSG = shared spiking graph, CBG = correlation based graph, and STTC across all 37 recordings encompassing high (n=10), moderate (n=14), and low (n=13) activity recordings. The Kruskal-Wallis test revealed significant differences for number of nodes and edges, the mean degree, network density, clustering coefficient and average shortest path length.

Notably, the number of nodes differed substantially (H = 19.6, p < 0.001) with CBG-method producing the lowest number of nodes (median = 72) compared to SSGs (median = 158, p = 0.0087, d = 0.71) and STTC-graphs (median = 181, p < 0.001, d = 1.14). This is naturally a result of the dual-threshold filtering for CBG-method where nodes must surpass the null-model significance test and the fixed correlation threshold. This results in low active nodes being excluded from the network.

Clustering coefficent exhibited the largest divergence between methods (H = 17.19, p < 0.001). SSGs showed significantly lower clustering (median = 0.48) compared to both, CBGs (median = 0.85, p < 0.001, d = 0.86) and STTC-graphs (median = 0.8, p = 0.027). STTC-graphs and CBGs did not differ significantly. The low clustering in SSGs is due to the z-score normalization that eliminates edges below the expected co-activation rate, resulting in sparser graphs.

### 2. Activitiy-based Network differences are method specific

After establishing that the three observed methods construct different networks, we were interested how methods capture the differences between activity levels. We found that all three methods detect significant differences in number of nodes (Kruskal-Wallis Test, SSG: H = 19.91, p < 0.001; STTC: H = 20.78, p < 0.001, CBG: H = 6.59, p = 0.037) and number of edges (Kruskal-Wallis Test, SSG: H = 13.35, p = 0.0013; STTC: H = 13.31, p = 0.0013, CBG: H = 9.7487, p = 0.0076). For these metrics on relating to graph size, SSG and STTC showed large effect sizes (d = 2.26 – 2.62) for low vs. moderate and low vs. high (e.g., d = 2.55 for low vs. moderate SSG), while correlation showed a moderate effect (d = 1.18 for low vs. high). This is probably due to aggressive node pruning in the correlation based method resulting in a compressed range.

**Fig 4.**
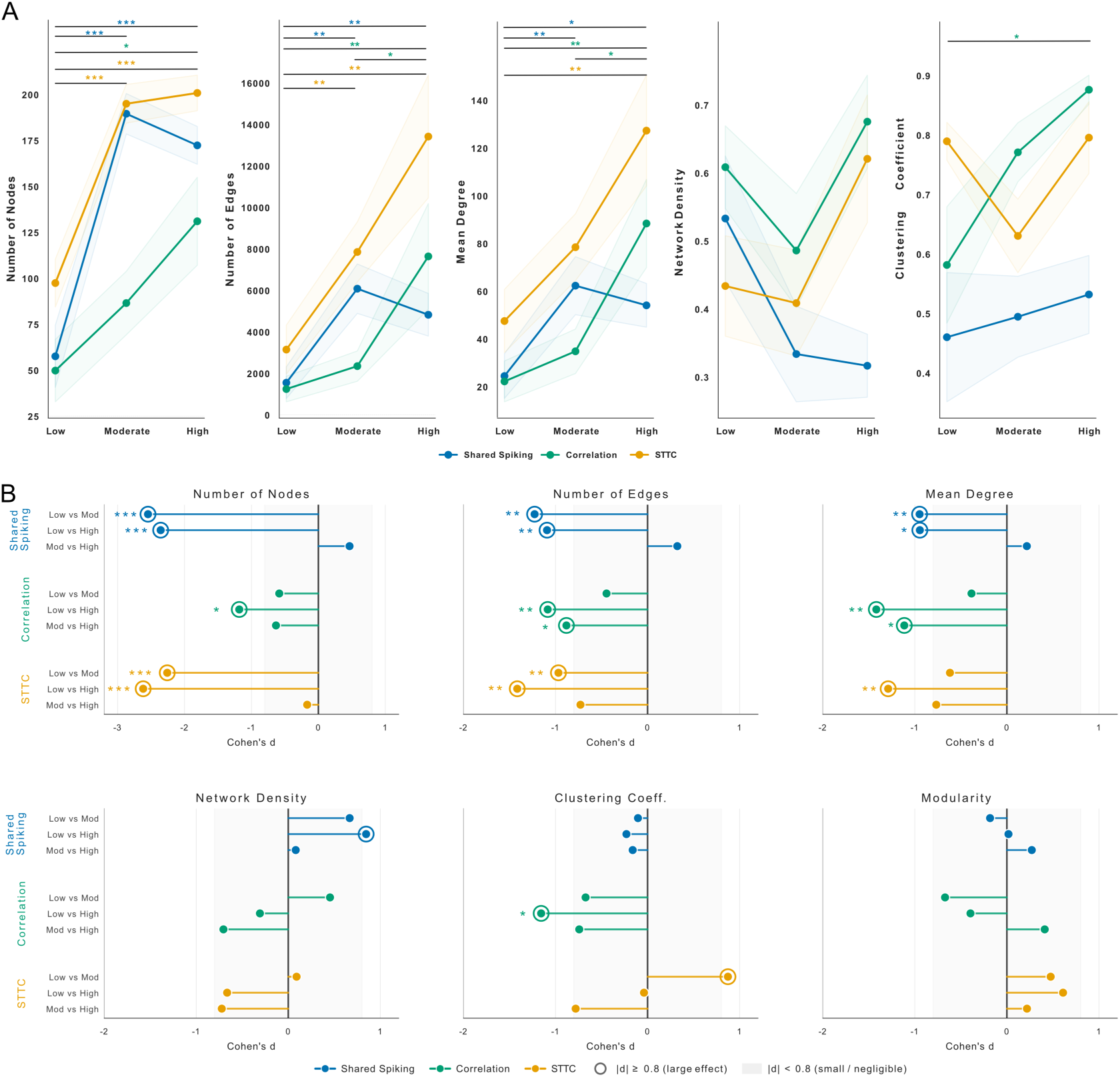
Method-specific sensitivity to neural activity level. **(A)** Mean (± SEM) network metric trajectories across low, moderate, and high activity recordings for each graph construction method. Coloured significance labels indicate pairwise test results (Dunn’s test) per method. All three methods detect activity-dependent increases in graph size (number of nodes, edges, mean degree), but diverge for topological metrics: clustering coefficient is activity-sensitive only for correlation-based graphs. **(B)** Pairwise effect sizes (Cohen’s d) for each activity group comparison, grouped by method. Lollipop length and direction indicate effect magnitude and sign; open circles mark large effects (|d| ≥ 0.8). The gray zone denotes small or negligible effects. Shared spiking and STTC show the strongest separation for graph size metrics (d > 2.0 for low vs. moderate/high), while correlation-based graphs show the strongest topological separation via clustering coefficient.

For topology metrics, methods diverged more clearly: Clustering coefficient was significant only for CGBs (H = 6.38, p = 0.041) which is a result of the large effect in low vs. high activity groups (d = 1.56, p = 0.014). This effect was not seen for SSG (H = 0.25, p = 0.88) or STTC (H = 4.67, p = 0.097). However, mean edge weight was activity sensitive for STTC (H = 8.79, p = 0.012) and SSG (H = 10.66, p = 0.005) but not for CBG (H = 4.94, p = 0.085). This is biologically meaningful since correlation weights are bounded by the coefficient range and thus show less sensitivity to activity-driven changes in synchronization.

While network density did not reach any significance, it is interesting to see that density was lower for moderate activity recordings compared to low activity recordings. More active recordings do not automatically yield denser graphs because the number of nodes can increase more than the relative number of edges. This is an important observation when reporting network metrics.

### 3. Normalization methods beats bin size in Shared Spiking Parameter Sensitivity

**Fig 5.**
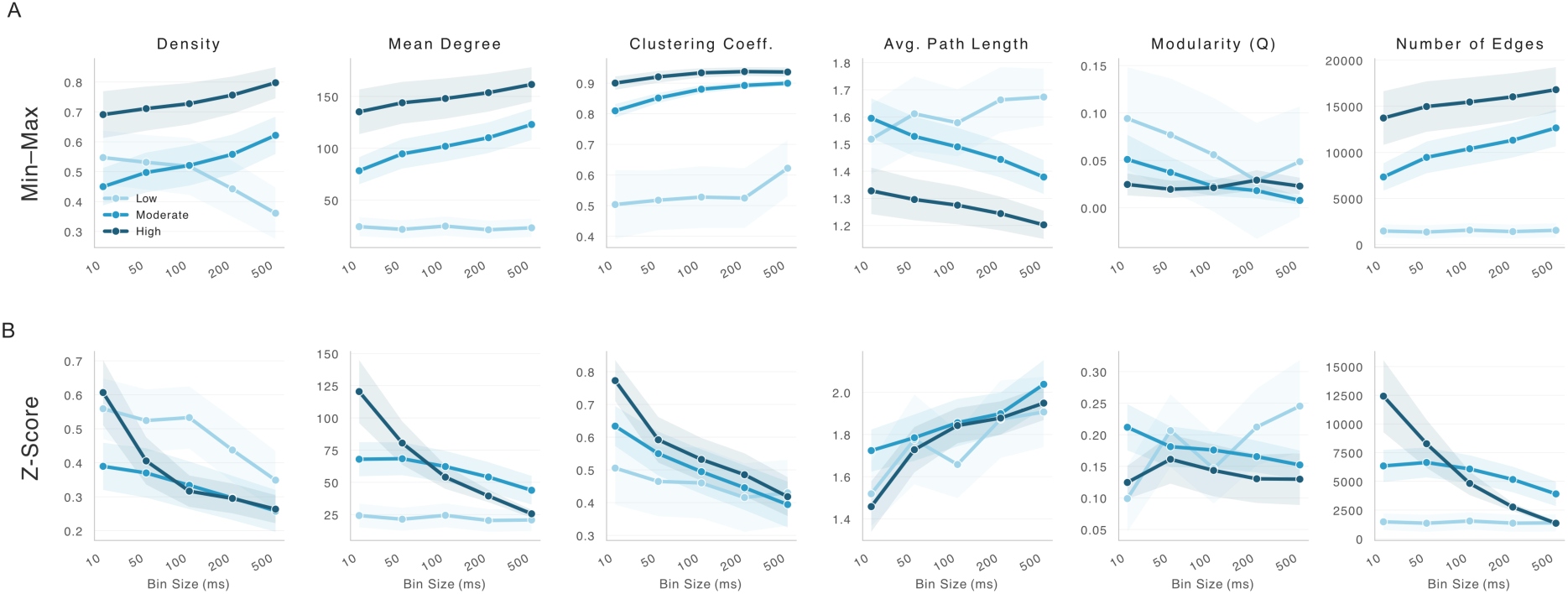
Parameter sensitivity of shared spiking graph construction. Mean (± SEM) network metrics across five bin sizes (10–500 ms) for **(A)** min–max normalization and **(B)** z-score normalization, stratified by activity level. Non-z-score normalizations (including min–max) preserve the edge set and produce nearly identical topology metrics, differing only in mean edge weight. Z-score normalization fundamentally alters graph structure by removing edges below the expected co-activation baseline, yielding sparser networks with lower clustering, longer path lengths, and higher modularity. Bin size exerts secondary effects, primarily increasing edge count and mean degree at coarser resolutions due to the higher probability of temporal co-occurrence within wider bins.

After investigating the impact of recording activity for each method, we wanted to explore the influence of parameter choice within each methods influences the resulting network structure. To asses this, we performed systematic sweeps for all constructing methods.

For SSG, we evaluated all 25 combinations of bin sizes (10, 50, 100, 200, 500ms) and four weight normalization methods next to the raw graphs (rate-normalized weights, co-activation proportion, standard min-max normalization, and Z-score normalization). The Kruskal-Wallis test showed significant differences for 7 of the 8 investigated metrics (all p < 0.001, except number of nodes: p = 0.72). However, these differences were dominated by normalization choice, while bin size had comparatively modest effects. For the non-Z-score metrics and raw graphs, the topology metrics were identical or nearly identical at each bin size (for density, clustering coefficient, average path length, and modularity, comparison between same bin size with different normalization pairs did not show any differences for all combinations of raw, rate-normalized, or min-max normalization at every bin size). This is explained by the way these normalizations work; In essence, they transform edge weights after graph construction which will preserve the edge set and thus preserving all the topology metrics that rely purely on network structure. Co-activation does preserve the edge set as well, but differs in mean edge weight due to the pairwise-specific scaling.

On contrast, Z-score normalization produces different graphs because it centers the edge weights around the expected co-spiking count, thus it alters the edge set itself. This resulted in large effects across all topology metrics: clustering coefficient differed by d = 1.53 between 500 ms-none and 500 ms-z_score (p < 0.001), density by d = 1.06 (p < 0.001), and modularity by d = 1.17 (p < 0.001). The z-score normalization consistently produced sparser networks (lower density, fewer edges), lower clustering, longer path lengths, and higher modularity compared to all other normalizations at the same bin size.

The bin size effects self were secondary compared to normalization methods. increasing bin size from 10 ms to 500 ms produced moderate shifts in path length (d = 0.88, p < 0.001) and clustering (d = 0.73, p = 0.002), reflecting the broader temporal connection window. Within non-z-score normalizations, bin size effects on density and clustering were negligible (d < 0.23), though mean degree and edge count increased with bin size as expected from the increasing probability of temporal co-occurrence at coarser resolutions.

### 4. Correlation Threshold Modulates Network Sparsity

**Fig 6.**
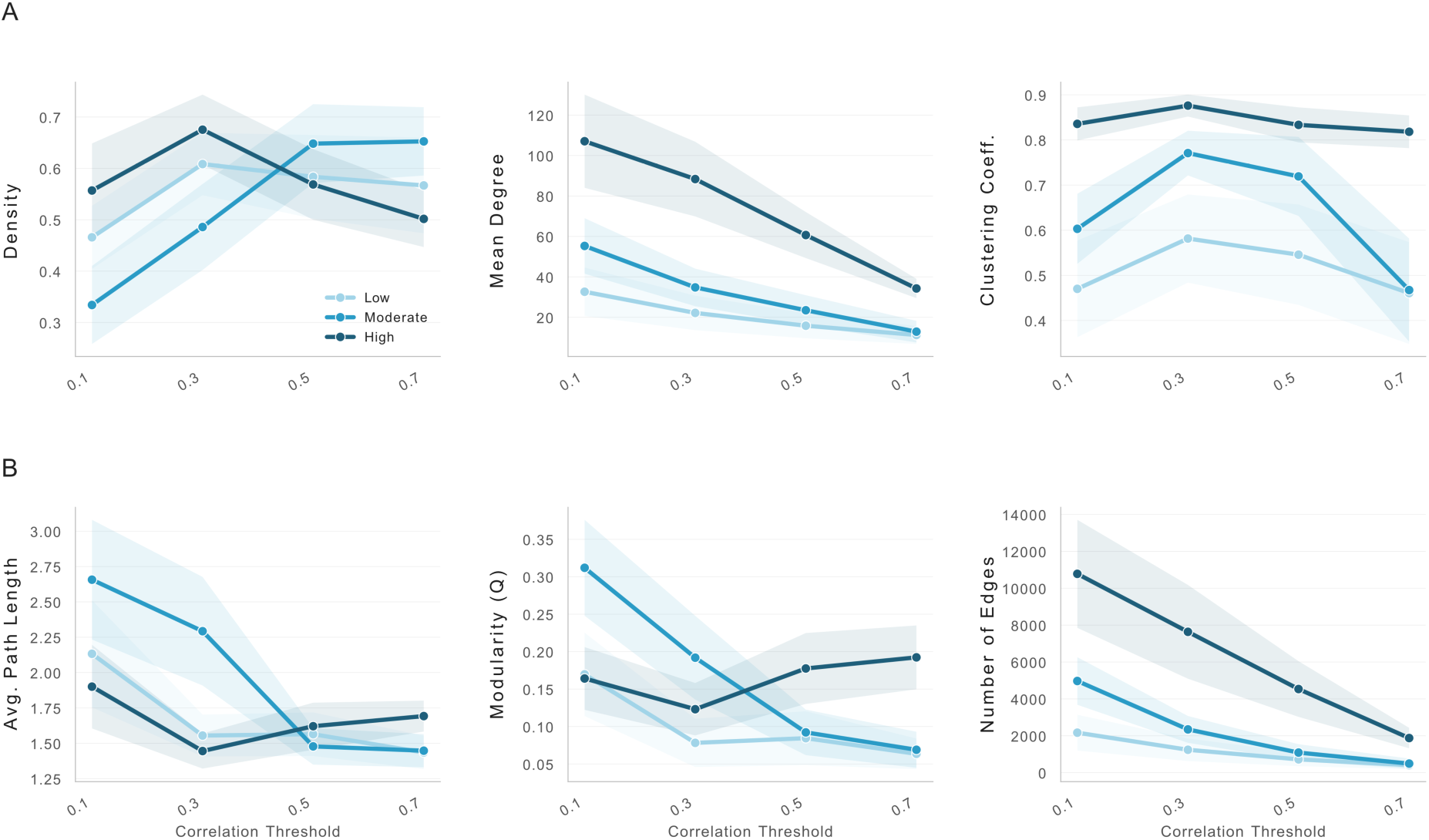
Parameter sensitivity of correlation-based graph construction. Mean (± SEM) network metrics across four correlation thresholds (0.1–0.7) applied after null-model significance testing, stratified by activity level. **(A)** Increasing the threshold progressively reduces mean degree and edge count as only the strongest correlations survive, while density and clustering coefficient remain relatively stable. **(B)** Average path length decreases at intermediate thresholds as peripheral low-degree nodes are pruned and modularity declines as community boundary nodes are lost.

We investigated the correlation method across four fixed thresholds (0.1, 0.3, 0.5, and 0.7) applied after null-model significance testing. Increasing the threshold reduced the number of nodes (given all pairwise comparisons for 0.1 vs. 0.7 for all activity levels, between 0.1: median = 138 and 0.7: median = 13, p < 0.001, d = 1.22), the number of edges (0.1: median = 2497, 0.7: median = 41, p < 0.001, d = 0.97), and the mean degree (0.1: median = 36.95, 0.7: median = 6.31, p <0.001, d = 0.95) while the mean edge weight increased (0.1: median = 0.35, 0.7: median 0.8, p < 0.001, d = 2.51) since with the higher threshold only strong correlations survive. Notably, the edge weight is defined by the correlation in our approach. Modularity decreases with stricter threshold (0.1: median = 0.13, 0.7: median = 0.08, p < 0.005, d = 0.71) representing the loss of peripheral nodes that normally form community boundaries. Density and clustering coefficient were not significantly affected through threshold ranges (density: H = 7.70, p = 0.053; clustering: H = 5.14, p = 0.16). The separation between activity groups was maintained across all tested thresholds (Cohen’s d for the low-versus-high contrast in mean degree remained above 1.2 at every threshold (range: 1.24–1.50), and for number of edges above 1.08 (range: 1.08–1.23). This suggests that the method’s ability to differentiate activity levels is more or less robust to reasonable threshold choices, even if the absolute network size can change substantially.

### 5. STTC exhibits intrinsic parameter robustness

We compared three tiling windows (at 10, 25, and 50ms) for the STTC method. In contrast to all other methods, no metric showed a significant difference between lag windows (Kruskal-Wallis for all at width p > 0.29). Also, pairwise comparisons showed small effect sizes with the largest being d=0.4 for mean weight between 10 and 50 ms (p = 0.13). The number of nodes, edges, and density were virtually unchanged across windows. Notably, the STTC method maintained stable separation across activity groups with Cohen’s d for low vs. high activity groups mean degree between 1.29 to 1.32 for the tiling windows and for number of edges from 1.40 to 1.42 (compare Fig. 7). This combination of relative parameter robustness and preserved biological sensitivity stands out for the STTC technique.

**Fig 7.**
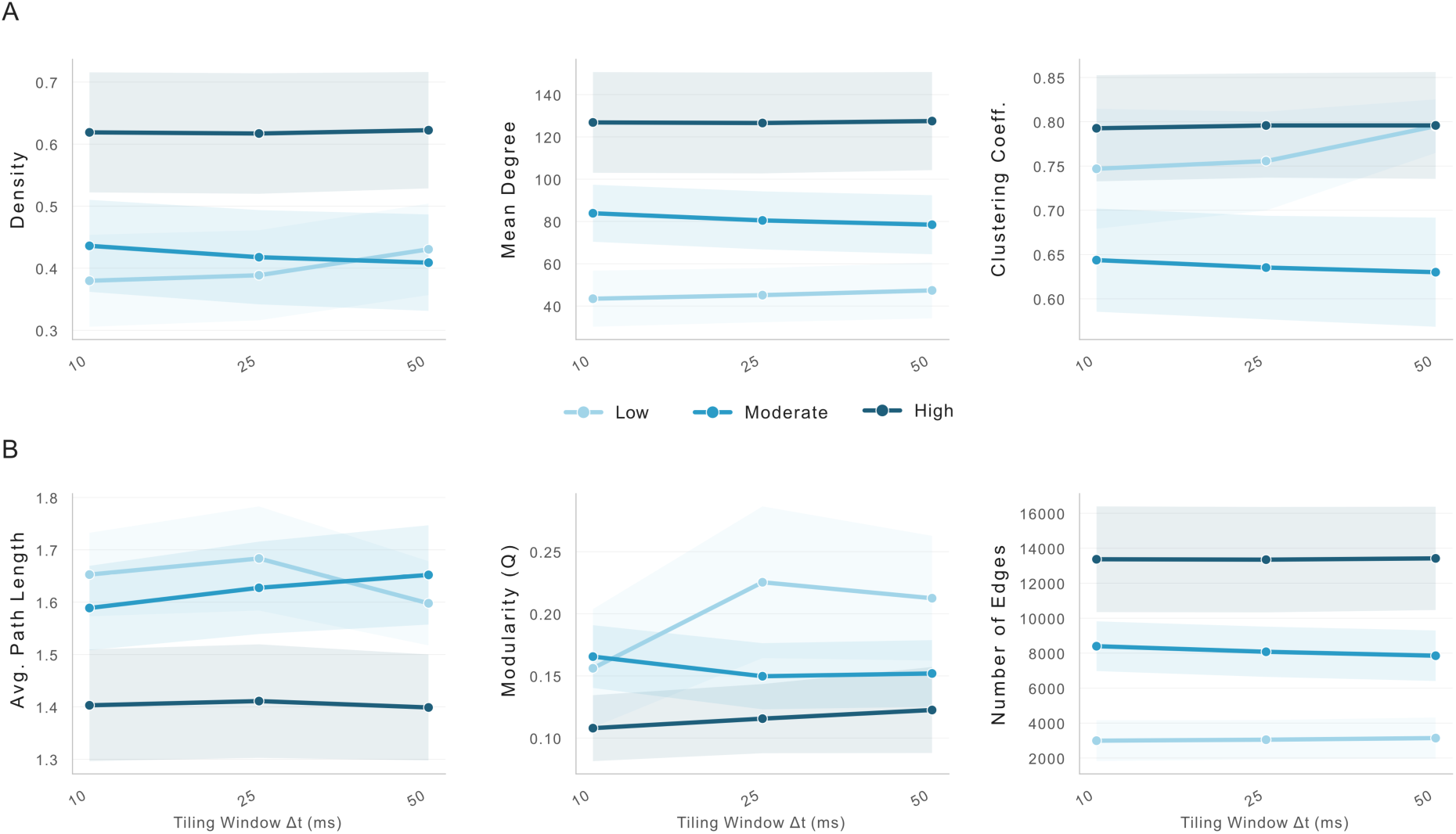
Parameter sensitivity of STTC graph construction. Mean (± SEM) network metrics across three tiling windows (Δt = 10, 25, 50 ms), stratified by activity level. **(A)** Density, mean degree, and clustering coefficient show minimal variation across window sizes. **(B)** Path length, modularity, and edge count are similarly stable. No metric differed significantly between tiling windows (all Kruskal-Wallis p > 0.29), with the largest pairwise effect being d = 0.4.

#### Edge Weight distributions and degree topology reflect method specific semantics

We plotted complementary cumulative distributions functions (CCDF) plots in a log-log space. Here, SSG exhibited the broadest degree range, while CBG and STTC showed steeper CCDF tails corresponding with their higher density. In general, high activity recordings had a narrower, more right-shifted degree distribution compared to low-activity recordings, reflecting the transition form sparser networks to more dense and synchronized connectivity topologies (Fig. 8 A).

**Fig. 8.**
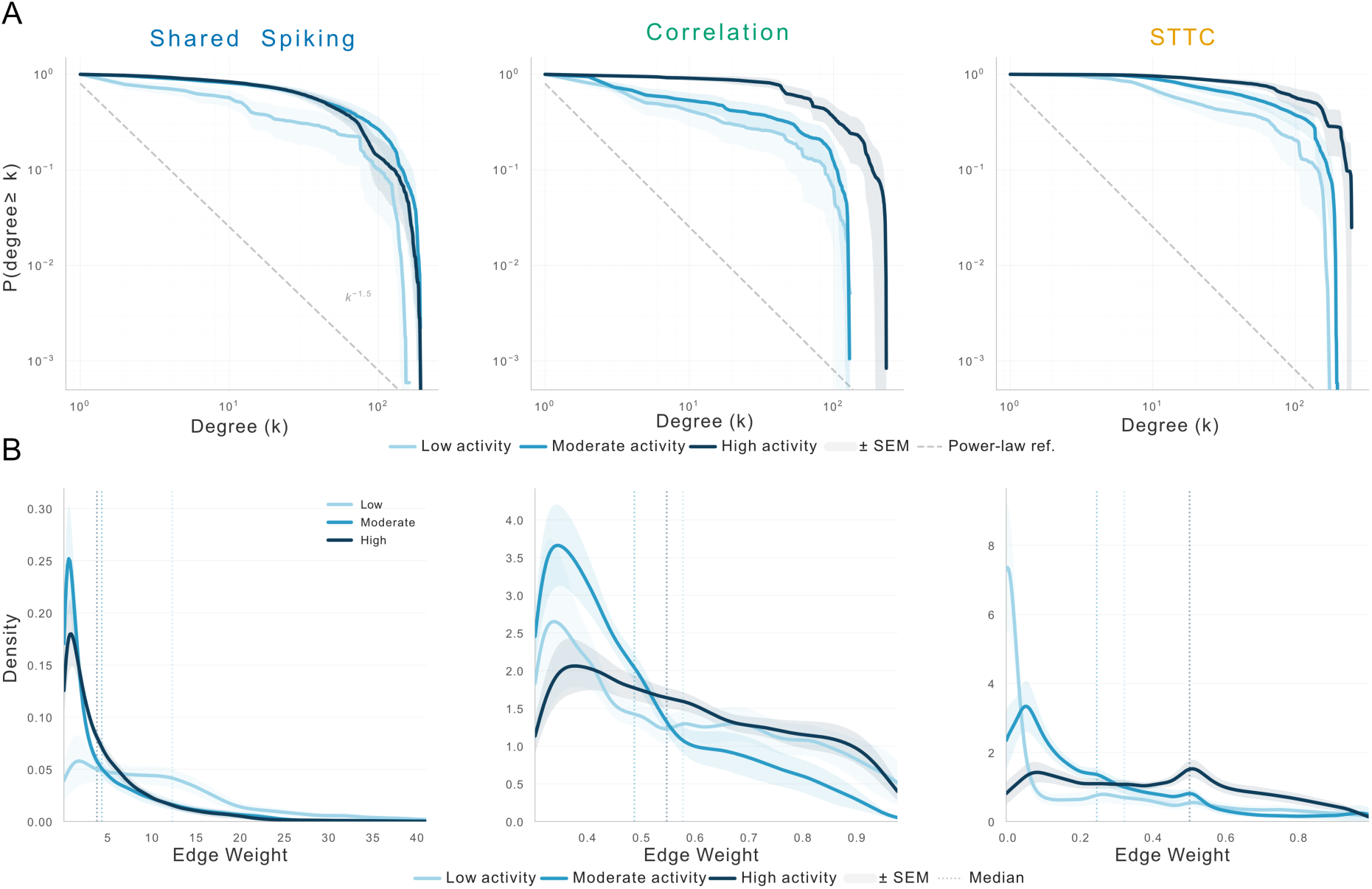
Degree and edge weight distributions across graph construction methods. **(A)** Complementary cumulative distribution functions (CCDF) of node degree on log–log axes (mean ± SEM across recordings). Dashed line indicates a power-law reference (k⁻¹·⁵). Shared spiking produces the broadest degree range, while correlation and STTC show steeper tails consistent with their higher network density. **(B)** Edge weight density distributions (mean KDE ± SEM). Dotted lines mark group medians. Shared spiking weights are heavily right-skewed, correlation weights are uni-modal and bounded by the coefficient range, and STTC weights peak near zero for low-activity recordings but broaden with increasing activity. The distinct distributional shapes confirm that the three methods impose fundamentally different weight structures on the same underlying spike data.

Edge weight distribution show further fundamental differences across methods. The SSG method produced a highly right-skewed distribution with a long tail representing highly synchronous electrodes. Meanwhile, CBG produced uni-modal distributions with a small rightward shift increasing with activity. The distributions is of course influenced by the chosen cut-off 0.3. STTC showed a peak near zero for low activity recordings while presenting a rather broad distribution with more uniform shape for high activity. Individual recording shapes (Fig. 8 B) showed similar shapes within their groups. This confirmes that these patterns are group-level features and not mere artifacts.

### 6. Inter-method agreement improves with activity level

Lastly, we quantified the overlap between the constructed networks by the different methods on the same recordings using the Jaccard index (fraction of shared edges) in relation to the respective firing rate of each recording using the Pearson correlation.

Interestingly, edge overlap was low across all method pairs (Fig. 9). For low-activity recordings, median Jaccard index ranged from 0.08 (SSG vs. STTC) to 0.20 (SSG vs. CBG) with substantial inter-recordings variability. The agreements improves with activity showing a Jaccard index of 0.13 (SSG vs. STTC), and 0.45 (CBG vs. STTC) for high activity recordings. CBG and STTC showed the highest overlap. This is likely explained because both methods involve a pairwise statistical testing (the null-model thresholding for CBG and the surrogate-based for the STTC). Pearson correlation for each method pair compared to the firing rate showed low correlation for SSG vs. CBG (r = −0.18, p = 0.274) and vs. STTC (r = −0.04, p = 0.833). Only CBG and STTC show a convincing correlation with statistical significance (r = + 0.45, p = 0.006) (compare Fig. 9 A).

**Fig 9.**
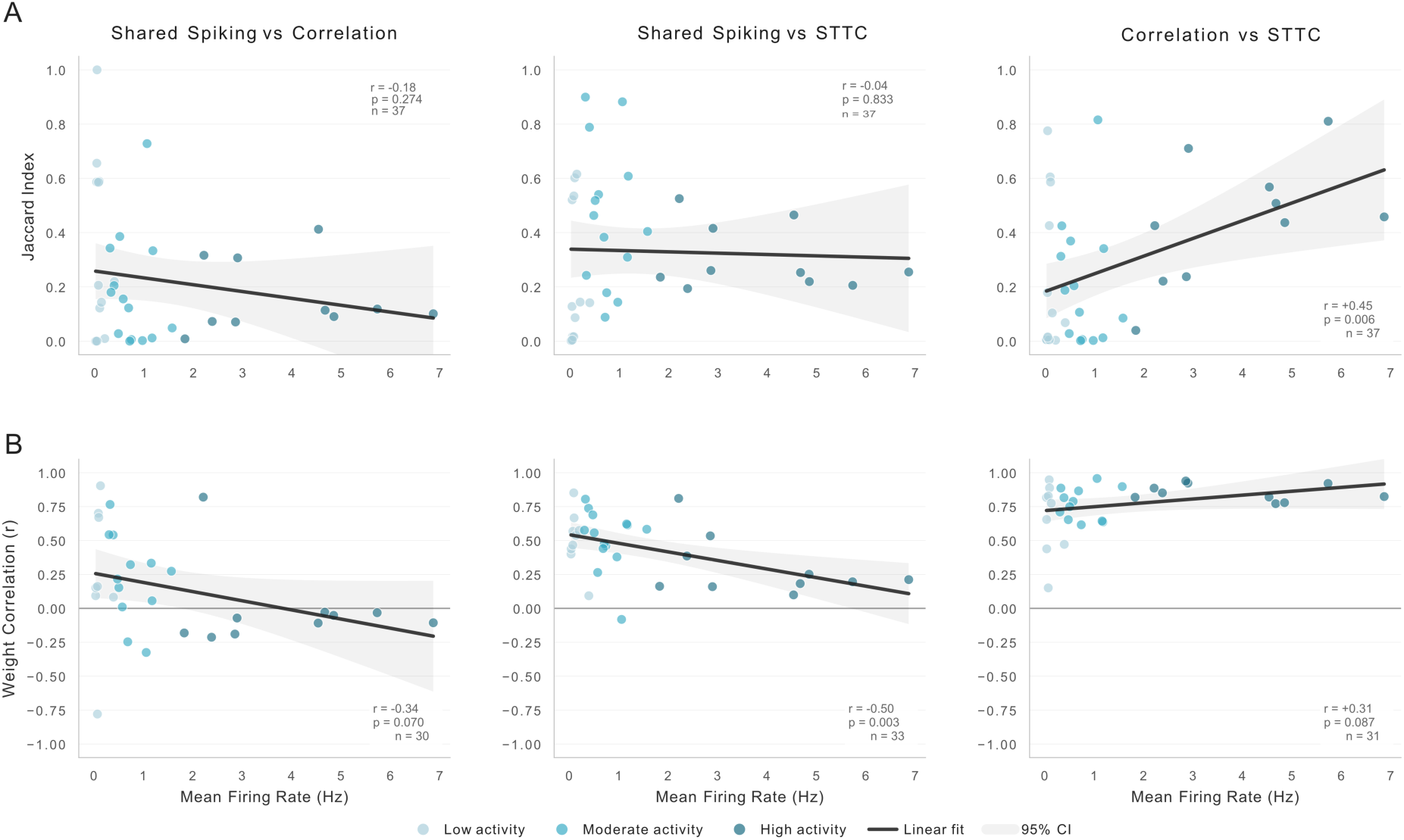
**(A)** For each recordings the Jaccard index is plotted against the firing rate and shows the fraction of shared edges for each pair of construction methods. Inter-method agreement is low but shows a positive correlation (r = 0.45, p = 0.006) for CBG and STTC with better agreement for higher activity. (B) Plotting the edge weight correlation for agreed edges shows negative correlation for SSG compared to CBG (r = −0.34, p = 0.07) and STTC (r = −0.5, p = 0.003), while CBG and STTC show a relative agreement. This highlights the fact that even if methods agree on graphs, they not necessarily agree on the importance of the edges.

Weight correlations on shared edges correlated to the recordings firing rate showed similar results (Fig.9 B) differed from these findings. CBG and STTC showed strong positive weight correlation increasing with activity (low activity: median r = 0.63, moderate activity: median r = 0.76, high activity: median r = 0.85), indicating that if these two methods agree on the existence of an edge, they also agree on its strength. Correlating this finding with firing rate we found a positive correlation (r = 0.31, p = 0.087), albeit without statistical significance.

SSG and STTC showed moderate positive correlation in low and moderate activity (median r = 0.50 – 0.55), but then declined for high activity (median r =0.22). Notably, the SSG and CBG showed progressive inversion with activity with a weak positive correlation for low activity recordings (median r = 0.22, with a high spread of correlations) and negative in high activity (median r = −0.08). This means that in highly active recordings SSG and CBG assign somewhat opposing weight rankings to the same edges. This could be explained by the z-score normalization that penalizes high-firing electrodes. This effect is also results in negative correlations for all recordings in relation to the firing rate this results in a negative correlation for SSG vs CBG (r = −0.34, p = 0.070) and SSG vs STTC (r = −0.50, p = 0.003).

Our results demonstrate that graph construction method choice introduces systematic and quantifiable differences in network topology that interact with the biological signal of interest. The three methods agree on higher-order organization (modularity, path length) but diverge in local structure, with each method showing a distinct sensitivity profile to activity-level differences. Parameter choices within methods matter unevenly: z-score normalization in shared spiking and threshold choice in correlation each alter topology substantially, while STTC is effectively parameter-free across the tested range. The low inter-method edge overlap, even in high-activity recordings, underscores that these methods are not interchangeable but construct fundamentally different representations of the same underlying neural activity.

## Discussion

Our central finding is that graph construction method choice results in systematic, quantifiable differences in network topology that can exceed the biological effect that is examined for. This finding is in line with the available body of the literature showing the dependence of estimated connectivity on the leveraged algorithms.[27–30] However, we extend on this body with to our knowledge the first publication benchmarking different methods using graph theory metrics on real world human cortical slice culture recordings. While the methods agreed on higher-order features including path-length or modularity, we found significant differences in local structure, biological sensitivity depending on slice activity, and the intra-method parameter dependence. These results have major implications for reproducability and interpretability of MEA-based network analysis, especially considering the variability of human tissue recordings.[1,2]

### Influence of parameter choices

We demonstrated that parameter choice can heavily influence the topology of the resulting graph. Depending on the method, normalization and thresholding are particularly influential on results with some parameters being shifted by Cohen’s d > 1.5 on the same recording.

Classically, correlation of pairwise connectivity estimations are confounded by the firing rate making some form of normalization necessary.[8,31] For our shared spiking approach, the z-score normalization is mechanistically different from the other normalization methods, because it redefines what does count as edge, while the others only rescale the edge weight. This also results in the counter-intuitive observation that with increased bin size, mean degree and number of edges decrease. In the non z-score normalization methods, bigger bin size means more temporal overlap and thus also increased graph size. However, for z-score normalization, larger bins also increase the expected co-activation that rises even faster for high firing nodes. This means that for frequently active electrode pairs at coarser temporal resolution leading to filtering edges that would survive at short bin size. Z-score normalization implicitly corrects for firing rate confounds making it conceptually closer to a null-model (like in STTC) [8]. The z-score normalization assumes a Poisson-like distribution model. However, in global network events, the co-activation does not stem from a pairwise coupling but a community synchronization. This positions it as the methodologically most principled choice within the shared spiking family. However, this choice comes with the cost of reduced sensitivity to global synchronization pattern that themselves are of not only biologically meaningful but also highly influence a recordings firing rate [32,33]. This has practical consequences since the choice of normalization method has larger effects on the networks topology (Cohen’s d = 1.07 – 1.53 for network density, clustering or modularity) than bin size for within the normalization (d < 0.23 for non-z-score methods, d = 0.45 – 0.88 for z-score across the full 10-500ms range).

We also want to highlight the importance of threshold wich still is an open question in neuronal connectivity interference across methods.[34–37] In our methods, CBG was most dependent on threshold choice since this was the primary criterion for edge acceptance, while thresholding was indirect in other methods using different time windows. However, all methods can theoretically be altered using additional thresholds, as for example a minimum edge weight after normalizing a shared spiking graph. We saw the massive influence these direct and indirect thresholds have on the investigated network parameters (compare Figure 5 and 6). We strongly recommend consequent reporting of any chosen thresholds, and advocate for further future work benchmarking different and adaptive thresholds comparable to suggestions in other modalities such as fMRI.[36,38,39]

### Method-specific sensitivity profiles

While STTC is the most robust method across all activity levels with the best parameter robustness (Kruskal Wallis p > 0.29, maximum Cohen’s d = 0.4), all three methods present specific sensitivity profiles. SSGs are more sensitive to node count and mean weight, especially when not normalized using the Z-Score method. With larger bin size, the probability of a spike-coactivation is higher for a relatively inactive node. This will result in increased node count, but a much lower network density (compare Fig. 5). This effect is highlighted with non-z-score normalizations. Indeed, there is a strong argument for this particular strength of the shared spiking approach since it also depicts weak (i.e., low active) nodes.[40] This might be an essential advantage in human tissue considering the variability in firing rate, even more so when used for pathomechanistic experiments.[2,41]

The correlation method showed significant differences for clustering coefficient. This arises from the method’s dural-threshold design with a null-model significance filter and a fixed correlation threshold. CBG selects edges for genuinely correlated activit and not coincidental co-firing. This effect will produce highly local dense neighborhoods.

STTC showed a combination of strong activity sensitivity (Cohen’s d = 1.29 – 2.62 for metrics across activity groups) with good parameter stability. Since the mathematical properties of STTC normalizes for firing rate, the additional circular-shift surrogate mechanic also adapts the threshold to each recording’s temporal structure.

### Inter-method agreement and the question of ground truth

Considering the low Jaccard indices for edge overlap across method pairs, we demonstrate that the three methods do not merely differ in scale or threshold but identify substantially different edge topologies for the same data. For moderate activity recordings, SSG and STTC share the most overlap, while Correlation and STTC had higher agreement in high activity slices. However, even for these method pairs, overlap was fewer than half of the available edges. This of courses raises the fundamental question which method captures the most “true” connectivity.

Since ground truth data would require simultaneous intracellular recordings, this question is out of the scope of this paper. However, the argument could go for a complementary approach with STTC as robust baseline method accompanied by a second method depending on the biological question. CBG selects pairs that show consistent temporal relationships throughout the whole recording, while SSG also includes sparsely spiking electrodes emphasizing specific pairwise coupling. Considering the weight correlation between SSG and CGB across activity levels, the sign reversal (r = + 0.22 for low activity, −0.08 in high activity) underscores this divergence: Z-score normalization penalizes edges between high firing nodes through the while the same sustained co-firing rate between electrodes might be rewarded. Thus, the same biological observation is assigned completely different edge weights.

### Recommendations

Based on our results, three recommendations can be formulated for network analysis of MEA recordings. First, all methodological choices for graph construction, including method, bin size, normalization, thresholds, etc., should be explicitly reported considering the heavy influence on topology. The current trend to more standardized workflows and incorporation into open-source frameworks are already a positive development and we are optimistic that the MEA research community will continue this trajectory.[42–44] Second, we consider STTC with a surrogate-based thresholding as a robust default choice when parameter sensitivity are of concern[8], supporting its use in the literature and also recent integration into MEA-NAP, which is an excellent tool providing graph theory analytical abilities to a broader user base.[44,45] In our real-world data example the method showed a negligible dependence on the lag window while maintaining solid biological sensitivity. Third, we suggest to validate methods across at least one additional method. This method can be chosen regarding the research question. Additionally, establishing a standard reference recording, essentially a kind of ‘white balance’ for graph methods, could help researchers intuitively understand the intricacies of their chosen analytical approach.

### Limitations

There are several limitations that should be addressed when interpreting our results. The data set comprised 37 recordings from human cortical slice cultures divided into three groups using an activity classification depending on firing rate and network burst frequency. Depending on the threshold choices, these activity groups composition be substantially different resulting in altered results. With human brain slice cultures becoming more prevalent in the literature due to refined methods, we believe it is important to stress the variability of biological activity of human brain slice recording. We here focus exclusively on human slice cultures, since this method is gaining rapid traction in neuroscience and “hot topic” subfields including cancer neuroscience. However, generalizability of our finding on organoids, stem cells, or rodent cultures as well as other MEA platforms remains to be established. Further, our analysis treated each electrode as node without spatial consideration like distance or known anatomical features. Both aspects are valuable for further research.

## Conclusion

Network analysis in MEA recordings is not a neutral measurement. The graph construction depends highly on method and parameter choice. We demonstrate how each method choice from graph construction, normalization, and thresholding each introduce systematic bias with effect sizes sometimes exceeding the biological systems that are investigaed. Z-score normalization fundamentally reshapes shared spiking networks by acting as implicit null-model. STTC provides the most parameter robust results across our test range exhibiting strong biological sensitivity. The overlap between methods remains below 50% proving that these methods are not interchangeable but rather complementary. We showed a quantitative framework and recommendations for parameter selection and reporting standard to improve reproducibility in MEA-based network neuroscience.

## Methods

### Slice Preparation

Preparation of the human brain slice cultures has been extensively described by our group [2,41]. In short, patients that underwent neurosurgical procedures for brain tumor or epilepsy donated entrance and sparse tissue after written informed consent and approval from the local ethics committee (EK-067-20). Tissue used in this publication was obtained between 2022 and 2025. Human cortex tissue is then placed in ice-cold carbogenated artificial cerebral spinal fluid (aCSF) and processed into 250µm thick slices using a vibratome. Slices are then cut into smaller pieces and placed on 30mm Millicell culture inserts with 0.4µm pores (Millipore). After one hour of intermediate HEPES media culturing, slices are eventually cultured with human CSF (hCSF) and stored in an incubator. MEA recordings can be performed at different time-points but are usually between *day-in-vitro* 4-14. For this publication, we reused formerly processed slices to describe and define the analytical methodology.

### Micro-electrode array recordings

Human brain slices were recorded using a 252-channel MEA with a 30µm electrode diameter and 200µm electrode spacing (USB-MEA 256-System, Multi Channel Systems MCS GmbH). For recording, slices are placed on the 16 x 16 MEA grid and a weighted harp (ALA-HSG MEA-5BD, Multi Channel Systems MCS GmbH) is positioned on the slice. During recording, slices are constantly perfused with carbogenated aCSF. After at least 30 minutes of equilibration, recordings are performed with a sampling rate of 10kHz or 25kHz respective to the experimental protocol. All data used in this publication is re-used from other experiments and selected on visual inspection of the rasterplot to ensure a diverse selection of slice activity and network activation patterns that represent the typical variability in human tissue. Recordings included into this benchmarking analysis thus were obtained under different conditions including spontaneous recordings (n=12), high potassium (8mM, n=10), with 5µM or 10µM gabazine (n=14), or 30µM acetylcholine (n=1).

### Data Processing, Spike Detection and Artefact removal

The Multi Channel Systems (MCS) MEA system uses a custom data format which is converted into HDF5 files using the MCS data manager software. All further analysis is performed using custom python code. Data is imported and bandpass filtered (150 – 4500 Hz) using a Butterworth filter (4th order). For spike detection, a mean absolute deviation (MAD) threshold is calculated for each channel at −5 * MAD. Signal crossings below the threshold are aligned to the nearest minimum in a 3ms window, including a dead time of 1ms to prevent repeated counting. Due to the amount of data volume of a single recording, we analyze our data in subparts of 120 seconds increasing compatibility with available RAM performance. Spiketimes are then stored and used for further analysis. Spikes can be detected through different methods including advanced spikesorting algorithms, e.g., based on template matching [3,42]. The threshold-based approach used here can undersample spiketimes, but cut-out waveforms show robust identification of real action potentials. To reduce artifacts, every recording was discretized into 10ms snippets. For any snippet with >200 active electrode, an artefact is assumed and the spikes are removed from the spike train.

Off note, for the data in this publication no spike sorting was performed since we concentrate on the network dynamics [46]. However, our displayed analysis is equally usable for spike trains after spike sorting.

### Data-driven classification of Activity Groups

To examine the influence of different activity states on graph construction, we aimed for three distinct groups of recordings exhibiting high synchronized activity, high to moderate activity, and low activity. Initial group classification was done by visual inspection of raster plots with n=37 slices identified divided into high active recordings with synchronized activity (n=17), high to moderate activity (n=10), and low activity slices (n=10).

To establish objective classification criteria, we performed unsupervised clustering analysis on eight quantitative features: mean firing rate, coefficient of variation of firing rate, percent active electrodes, network burst frequency, percent of spikes in network bursts, mean network burst duration, mean participation percentage, and mean burst frequency per minute. K-means clustering with Ward’s linkage hierarchical clustering and DBSCAN were applied to identify natural groupings in the data. Optimal cluster number was determined using the elbow method, silhouette analysis, Calinski-Harabasz index, and Davies-Bouldin index.

The clustering analysis revealed k=5 as the optimal number of clusters, but consolidated into three biologically interpretable groups for functional analysis. The data-driven approach identified that firing rate and network burst frequency were the primary discriminators of activity states, while other features showed high within-group variability.

The cutoff values for firing rate and network burst frequency were iteratively refined to achieve balanced group sizes (n=13 low, n=14 moderate, n=10 high activity) while maintaining biological interpretability and respecting the natural cluster boundaries identified in unsupervised analysis.

### Functional Connectivity Graph Construction

To investigate the impact of graph construction methodology, we connectivity based graphs for each recording in parallel using three different methods to infere connectivity between electrodes and using different parameters within each method. Each method results in a weighted graph G where active electrodes represent the nodes (V) and edges (E) are defined as inferred connectivity. Edges are weighted (W).

### Simultaneous Spiking Graph Construction

The most simplistic graph construction method is coupling of coincident spike events between electrode pairs. Spike trains were first discretized into binary spike count vectors by binning spike times into non-overlapping time bins of width. Five bin widths were systematically evaluated: 10, 50, 100, 200, 500 ms. For each pair of electrodes, simultaneous activity in any of the bins leads to creation of an edge between electrodes with the number of co-occurrences as the raw edge weight. A minimum of two co-active bins is required for edge creation. Pseudo-code for this graph is in the supplementary material (compare algorithm S1).

For edge weight normalization, four methods were compared next to *raw counts* of co-occurrences: *Rate-normalized weights* are the raw count divided by the time length of the recording in seconds, resulting in a rate of co-activity per second*. Co-activation proportion* divided the shared count by the number of active bins by the less active channel, returning a value between 0 and 1 for each electrode pair. This can result in high weights for relatively unactive channels. *Standard min-max normalization* rescales all edge weights fo the graph between 0 and 1 using (W - W_min_) / (W_max_ - W_min_) applied to the full set of edges for each recording. *Z-score normalization* uses an expected number of co-occurences under a Poisson model (expected coincidences = firing rate₁ × firing rate₂ × Δt × T, where Δt is the bin width, and T is the recording duration; the z-score is computed as (observed – expected) / √expected, retaining only positive values).

The full parameter calculation consisted of all five bin widths crossed with five normalization methods, resulting in 25 parameter combination per recording. For comparison between methods, a 100ms bin size with z-score normalization was used.

### Correlation Based Graph Construction

For CBGs, binned spike trains with 100ms resolution were used and converted to spike count vectors. For each electrode pair, the Pearson correlation coefficient was computed at zero lag. At this bin resolution, the 100 ms bin width itself serves as the temporal integration window. We derived an adaptive threshold form a null model: For each recording, we calculated a surrogate distribution of correlation coefficients by randomly permuting (200 iterations) the bin order of one spike train from a randomly selected electrode pair and recomputing the distribution. Thus, the spike count for each channel is preserved while deconstructing a possibly functional connection. Between the electrodes an edge is then created if the correlations exceeds two thresholds: A fixed Pearson r threshold (calculated for 0.1, 0.3, 0.5, and 0.7) and the 95th percentile of the null model. Compare also the algorithm S2 in the supplementary material.

### Spike Time Tiling Coefficinet Graph Construciton

STTC graphs are constructed using the Spike Time Tile Coefficient as a measure of spike train correlation between two electrodes that is robust to firing rate differences.[8] STTC values are calculated as described and can range from −1 to +1 with 1+ indicating perfect temporal correlation while 0 suggests independence.[8] To evaluate statistical significance per electrode pair, the circular shift surrogate procedure was used with 180 surrogate values generated by shifting one spike train by a random amount drawn from [recording-duration/4, 3xrecording-duration/4] (compare pseudo-code in supplementary material algorithm S3). Comparatively to the CBGs, an edge was regarded as significant if the computed STTC exceeded the 95th percentile of the surrogate distribution. The edge weight is then set to the observed STCC for significant pairs of electrodes. For sensitivity analysis, we observed thre tiling window sizes: 10, 25, and 50ms. For comparison between methods, we used the 50ms window size.

### Graph Post Processing

After edge construction, all graphs were restricted to their largest connected component to ensure that path-based metrics (e.g., average shortest path length) remain well-defined and to avoid inflation of modularity estimates by small disconnected subgraphs. This approach follows standard practice in network neuroscience [47].

### Graph Theory Measurements

A standardized set of graph-level metrics was computed for every graph regardless of construction method. Network density was defined as the ratio of observed edges to the maximum possible. Degree statistics (mean, standard deviation, maximum) were computed from the unweighted degree sequence. Edge weight statistics (mean, standard deviation, minimum, maximum) captured the distribution of inferred connection strengths. The weighted clustering coefficient was computed following Onnela et al. (2005).[48] Average shortest path length was computed on the largest connected component using unweighted hop distance. Community structure was identified using the greedy modularity optimization algorithm [49] as implemented in NetworkX [50], and the resulting partition was used to compute the modularity index *Q*. The fraction of nodes belonging to the largest connected component was recorded as a measure of overall network fragmentation

### Statistical Anaylsis

Statistical analysis proceeded along two tracks. Track A addressed cross-method comparison: for each network metric, differences across the three graph construction methods were tested using the Kruskal-Wallis H-test. Dunn’s post-hoc test with Bonferroni correction for multiple comparisons was applied to identify which method pairs differed. Pairwise Mann-Whitney U tests were computed for all method combinations, and effect sizes were quantified using Cohen’s *d* with pooled standard deviation. This analysis was performed both pooled across all recordings and stratified by activity group. Track B addressed parameter sensitivity: the same non-parametric testing (Kruskal-Wallis + Dunn’s post-hoc) was applied within each method across its respective parameter sweep levels. For shared spiking, this tested the effect of bin size and normalization method on all network metrics; for cross-correlation, the effect of threshold stringency; and for STTC, the effect of tiling window width. Effect sizes (Cohen’s d) were computed for all pairwise parameter comparisons and categorized as negligible (|d| < 0.2), small (0.2– 0.5), medium (0.5– 0.8), or large (> 0.8).

## Use of Generative AI

Parts of the code were developed with the help of a generative-AI assistant, Claude (Anthropic) (model versions: Claude Sonnet 3.5–5, Claude Opus 4.6–4.8).Claude was used to help: draft and review the analysis code (spike processing, functional-connectivity, graph construction, network-metric computation, plotting); conceptualize, review, and sanity-check the statistical-analysis approach; review the results; improve documentation and the language of accompanying text; draft and review the accompanying manuscript. All AI-assisted code was executed, tested, and verified by the main author against expected outputs. No figures or data were generated by AI image tools. All figures are produced programmatically from the recorded data, except the methodological abstract which was build using biorender.com. The main author reviewed all AI-assisted output and takes full responsibility for the contents of this manuscript and the code repository.

## Code and Data Availability

All analysis code is available at github.com/ jonasort/Graph_Theory_Benchmarking_in_human_brain_slices/releases/tag/Human_Brain. The data ist archived on Zenodo (DOI: https://doi.org/10.5281/zenodo.21283609).

## Author Contributions

Conceptualization: JO, DD, HK. Tissue Harvesting: HH, AH, DD, HC. Experiment Conduction: VW, AB, JH, AKR, JO. Coding: JO. Analysis: JO. Manuscript Writing: JO. Manuscript correction: All.

## Supporting Information

### Algorithm S1 Shared Spiking Graph

Connect electrodes that frequently fire within the same time bin.

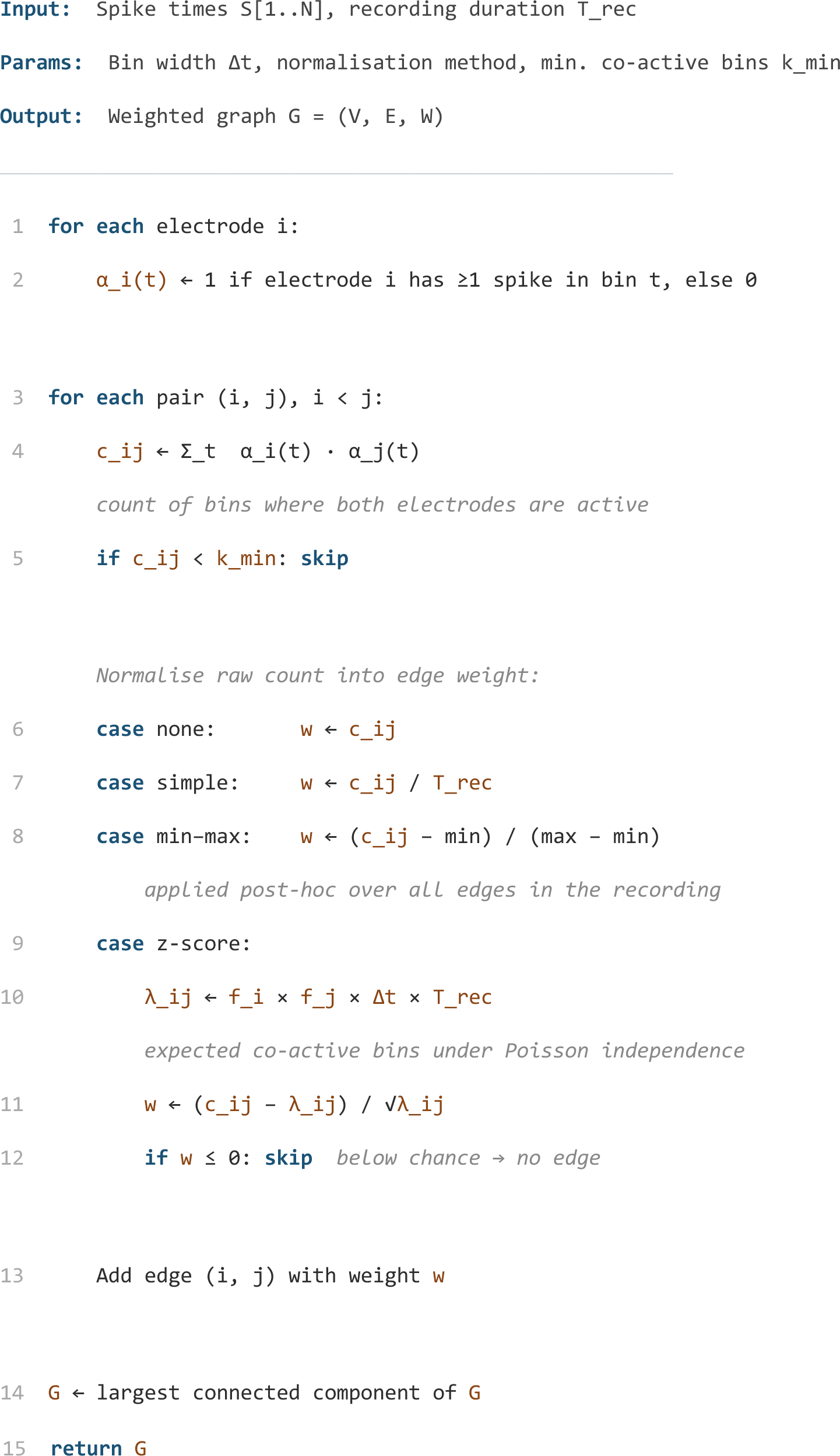

### Algorithm S2 Correlation-Based Graph

Connect electrodes whose firing rate patterns are correlated over time.

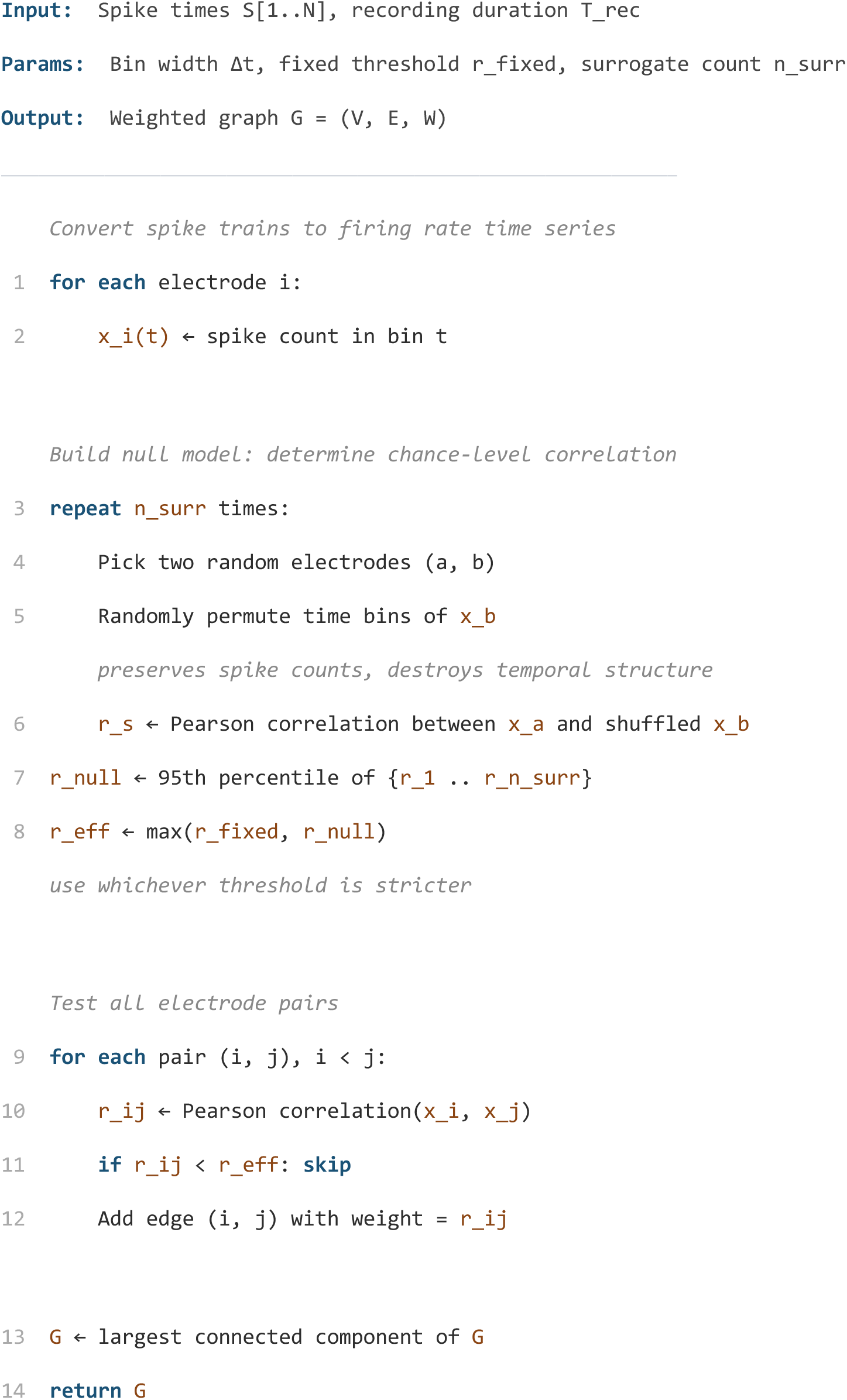

### Algorithm S3 Spike Time Tiling Coefficient (STTC)

Connect electrodes whose spikes occur near each other in time, more often than expected by chance. Operates directly on spike times without binning.

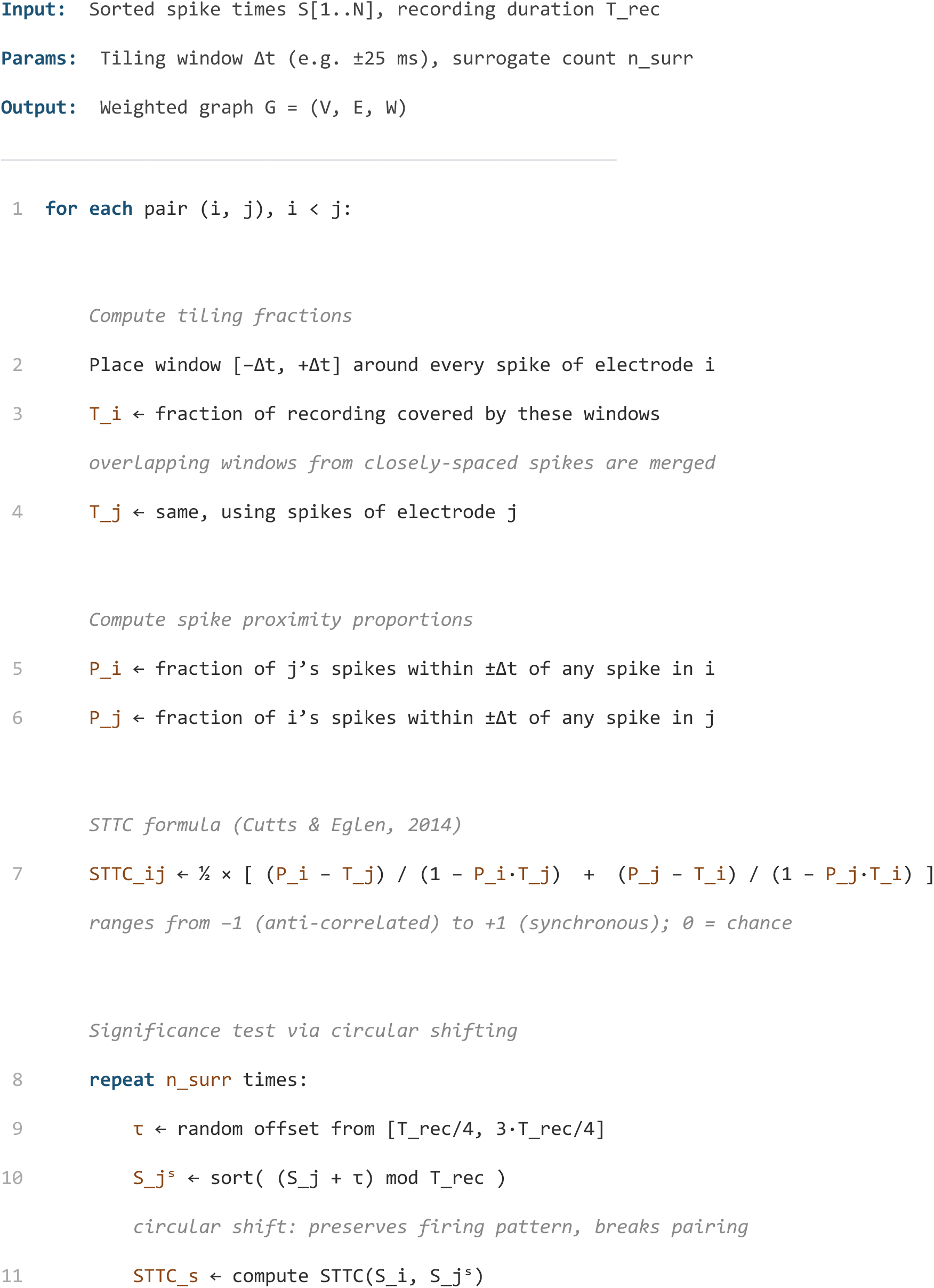
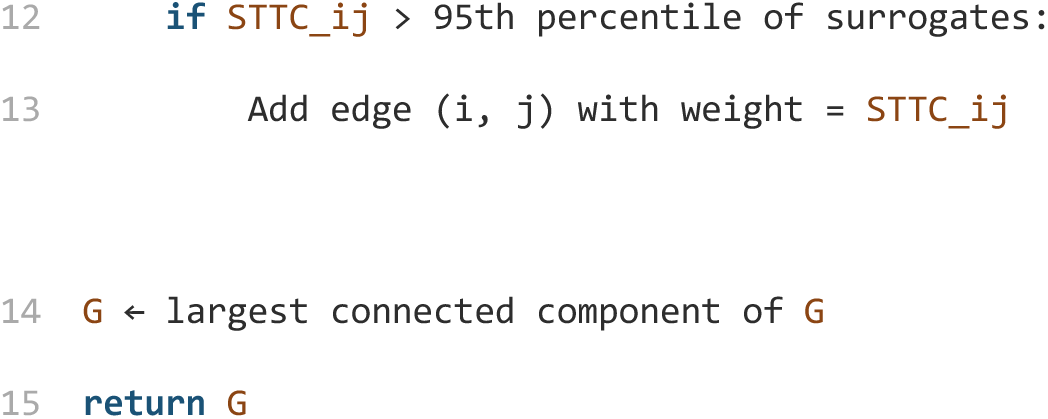

